# CausalTab: PSI-MITAB 2.8 updated format for signaling data representation and dissemination

**DOI:** 10.1101/385773

**Authors:** L. Perfetto, M.L. Acencio, G. Bradley, G. Cesareni, N.Del Toro, D. Fazekas, H. Hermjakob, T. Korcsmaros, M. Kuiper, A. Lægreid, P. Lo Surdo, R.C. Lovering, S. Orchard, P. Porras, PD. Thomas, V. Touré, J. Zobolas, L. Licata

## Abstract

Combining multiple layers of information underlying biological complexity into a structured framework, and in particular deciphering the molecular mechanisms behind cellular phenotypes, represent two challenges in systems biology. A key task is the formalisation of such information in models describing how biological entities interact to mediate the response to external and internal signals. Several databases with signaling information, such as SIGNOR, SignaLink and IntAct, focus on capturing, organising and displaying signaling interactions by representing them as binary, causal relationships between biological entities. The curation efforts that build these individual databases demand a concerted effort to ensure interoperability among resources, through the development of a standardized exchange format, ontologies and controlled vocabularies supporting the domain of causal interactions. Aware of the enormous benefits of standardization efforts in the molecular interaction research field, representatives of the signalling network community agreed to extend the PSI-MI controlled vocabulary to include additional terms representing aspects of causal interactions. Here, we present a common standard for the representation and dissemination of signaling information: the PSI Causal Interaction tabular format (CausalTAB) which is an extension of the existing PSI-MI tab-delimited format, now designated MITAB2.8. We define the new term “causal interaction”, and related child terms, which are children of the PSI-MI “molecular interaction” term. The new vocabulary terms in this extended PSI-MI format will enable systems biologists to model large-scale signaling networks more precisely and with higher coverage than before.

## Introduction

Cells are complex and dynamic systems responding to internal or environmental cues (Lodish et al., 2000). Chemical, physical or mechanical stimuli are sensed by receptor proteins which trigger the propagation, amplification and modulation of the signal through a cascade of enzymatic reactions and physical interactions, culminating in the rewiring of the gene expression profile (Lee & Yaffe, 2016). Collectively the intricate interaction mesh underlying these processes are referred to as “signal transduction”.

Given the importance that signal transduction has in determining cell phenotype, under either physiological or pathological conditions, obtaining a thorough understanding of the molecular mechanisms underlying the stimulus-phenotype relationships is one of the major goals of systems biology (Barabási & Oltvai, 2004).

To document and archive our growing understanding of signaling systems, a number of resources have undertaken the effort of retrieving from the scientific literature experimental observations supporting causal relationships between signaling proteins and to convert these into structured knowledge. These resources can be can be grouped according to the adopted data representation model in four main categories: activity flow, enzyme-substrate, indirect molecular interaction and process description (Türei *et al*, 2016).

Among the resources representing signaling information as activity flows, databases such as non-metabolic KEGG (Kanehisa *et al*, 2017), Reactome (Sidiropoulos *et al*, 2017), SIGNOR (Perfetto L *et al*, 2016) and SignaLink (Fazekas *et al*, 2013) focus on the capture, organization and display of signaling relationships and represent them as binary, causal relationships between biological entities. Databases such as IntAct (Orchard *et al*, 2014) focus on the capture of molecular physical interaction adding causality as an additional annotation of the interaction.

In addition, the Gene Ontology Consortium (GOC) (Ashburner *et al*, 2000) has long been providing causal statements in the form of annotations to terms from the gene ontology (GO) branches *regulation of molecular function* (GO:0065009) and *regulation of biological process* (GO:0050789). More recently, the GOC has implemented an extension of GO annotations called GO-Causal Activity Modeling (GO-CAM; http://geneontology.org/cam) (The Gene Ontology Consortium, 2017). GO-CAM provides a “grammar” for linking simple GO annotations into larger, semantically structured models (such as biological pathways) using relations from the Relations Ontology (RO) (Smith *et al*, 2005).

The emerging picture is of a fragmented and sparse collection of annotated signaling relationships, which is structured according to different curation approaches. As no single database is comprehensive, users face the challenge of integrating different datasets to obtain maximum coverage. To increase interoperability, adequate standards, ontologies and common export formats are required. The importance of producing standardized data for the life sciences has been documented many times. A particularly successful example of this is the work of the Molecular Interaction work group of the HUPO-Proteomics Standards Initiative (HUPO-PSI) (Deutsch *et al*, 2017). Over the last 15 years, this group has developed and disseminated community standards, tools and controlled vocabularies (CVs) (Kerrien *et al*, 2007) (Sivade Dumousseau *et al*, 2018) and minimum information guidelines for authors (Orchard *et al*, 2007). The ability to display molecular interactions in a single, unified PSI-MI XML format represented a milestone in the field of molecular interactions (Hermjakob *et al*, 2004), and the use of common controlled vocabularies has enabled the consistent annotation of the captured information. A tab-delimited format, MITAB, has proven more suitable for users requiring a simpler, human-readable configuration (Kerrien *et al*, 2007). Implementation of these standards by all the leading molecular interaction databases has considerably contributed to data exchange, representation and comparison and encouraged the development of specific tools, such as PSICQUIC (Aranda *et al*, 2011) and Cytoscape (Smoot *et al*, 2011) to retrieve and visualize this information.

XML formats handling signaling information are already available (Hucka *et al*, 2003). They support the representation of molecular interaction data using a reaction-based model that can store very detailed information about the physical and chemical properties of an interaction, such as equilibrium constant, reaction speed, reactant concentrations etc. Key reaction-based file formats are SBML (Hucka *et al*, 2003) and BioPax (Demir *et al*, 2010), which are more suitable for storing chemical interactions rather than molecular interaction networks.

However, curation into these formats carries a large resource overhead, management requires programming skill and the files are not readily human-readable. Experience from the PSI-MI workgroup taught us that MITAB is much more popular than PSI-MI XML, albeit much less expressive. During an EMBL/EC-funded Causal Reasoning Workshop organised at EMBL-EBI in 2016, representatives of some of the main signalling repositories (SIGNOR, IntAct (Causal Reasoning), GO, SignaLink 2.0) recognized the necessity to move toward unification and standardization of signalling data and therefore to adopt common standards and controlled vocabularies. At this meeting, it was agreed to adopt and expand the existing PSI-MI CV by adding new terms required to define different characteristics of causal interactions and to address the lack of a tab-delimited representation of causality by adopting and extending the CausalTAB format originally designed by the SIGNOR database curators.

We report here the first version of this common standard for the representation and dissemination of signalling information, to capture causal interactions among biological entities: the PSI Causal Interaction format TAB, developed in compliance with the PSI framework and adopted by the MI workgroup of the HUPO-PSI as an extension of the existing MITAB format (CausalTAB, MITAB2.8). We have defined a new controlled vocabulary root term, “causal interaction”, and its related child terms, to annotate the different aspects of causal interactions.

## Results

### Causal Interactions in the PSI-MI framework

CausalTAB is primarily inspired by the HUPO-PSI MITAB and has been designed to be PSI MITAB compliant, an Excel-compatible, tab-delimited format developed for users requiring only minimal information in a user-friendly structure. Since its development, the MITAB format has been extended to allow a more granular representation of the interaction data, with MITAB 2.6 and 2.7 versions now available (del-Toro *et al*, 2013). The data is described in the format using the PSI-MI CV developed to standardize interaction data and enable a systematic annotation of data. The PSI-MI CV has a hierarchical structure where each term can be mapped to its parent and child terms (Fig.1) and allows biocurators to consistently capture molecular interaction data and empowers users to perform systematically data searches.

The PSI-MI CV and MITAB were originally developed to capture a standardized representation of physical interactions; interaction directionality and the resulting effects (activation/inhibition for protein-protein interactions and up-/down-regulation for regulatory interactions) were not included. However, a large number of physical interactions are known to be regulatory, pointing to the need for extending the PSI-MI standard to incorporate additional layers of abstraction. CausalTAB/MITAB2.8 has therefore been designed to meet the needs of those members of the molecular interaction community who wish to extend their representation of molecular interaction data with directionality data.

### Revision and extension of PSI-MI CV and MITAB file structure into CausalTAB standard

In a biological context, causal interactions are abstractions representing the regulatory effect that a regulator entity (e.g, a stimulus, a transcription factor, an enzyme, etc) has on a target entity (e.g, a receptor, target gene, a substrate, etc). At their most basic level, causal interactions involve two partners, they have a direction (subject -> object) and a description of the regulatory effect (positive or negative). Chains of causal interactions underlie signal transduction, which modulates the cell response.

In order to enable the representation of causal relationships in a PSI-MI compatible format, we set out to scrutinize the data structure of resources annotating causal relationships to identify the need for extension of the PSI-MI CV. Given the binary, directed and effective nature of causal interactions, the process has been conceptually organised into four steps: 1) revision of terms to describe the directionality of the interaction; 2) revision of terms to describe the causality of the interaction; 3) revision of terms to describe the molecular mechanisms underlying the interaction; 4) revision of terms to describe the entities.

To this end, we first defined the new term ‘causal interaction’ as a new branch term in the molecular interaction CV (Fig. 1), and subsequently proceeded with the creation of related child terms. During this process, every new term was linked to a definition and a reference and was systematically integrated into the CV hierarchy. The ‘causal interaction’ term is defined as ‘Binary causative relationships between biological entities’. CV terms belonging to this term allow the description of causal interactions using the current PSI-MI schema. ‘Causal interaction’ is the parent of a set of other terms related to each other by hierarchical configuration (Fig. 1).

**Figure 1:**
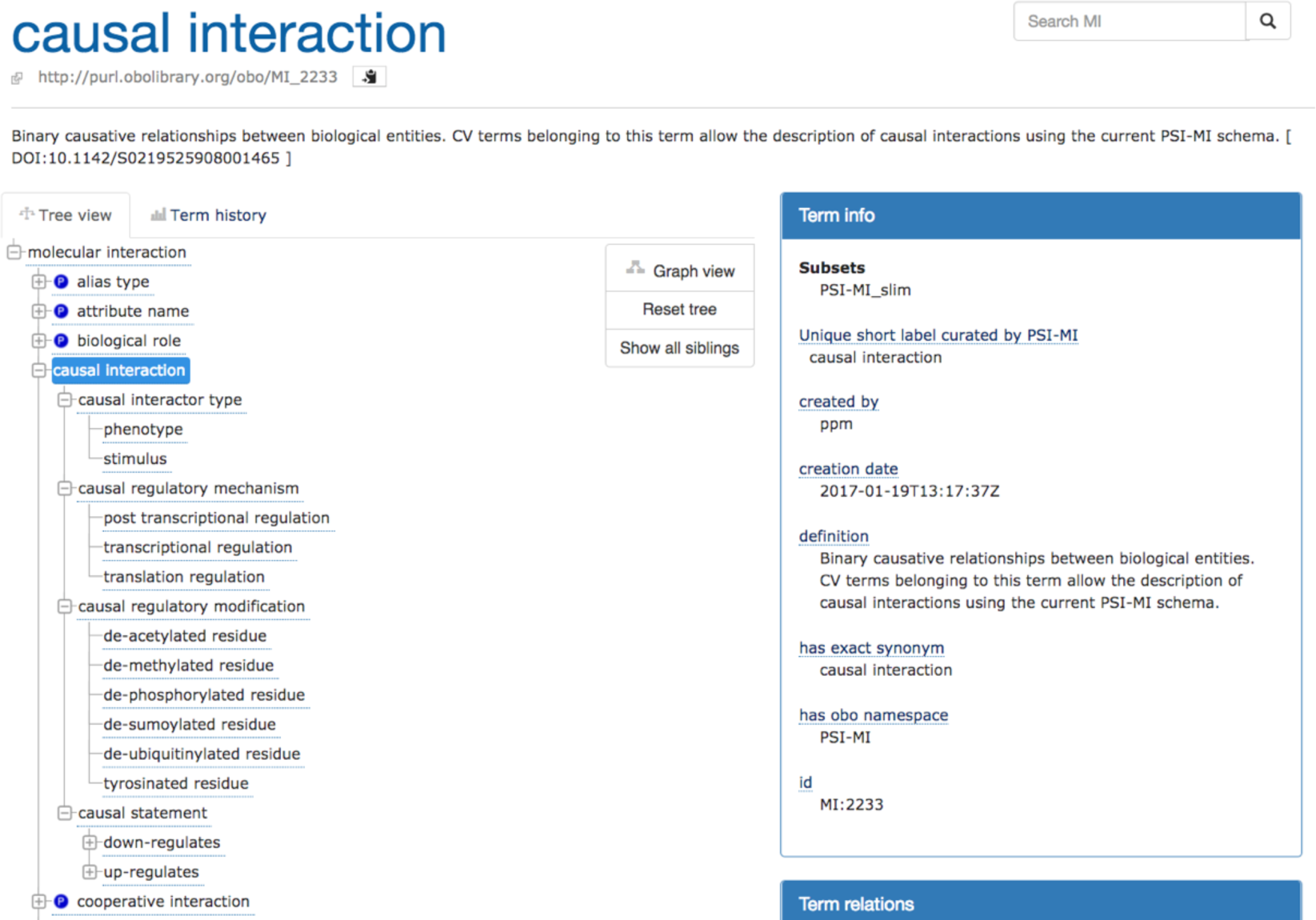
The hierarchical structure of the new PSI-MI controlled vocabularies terms. Causal Interaction root and children terms as shown in the Ontology Lookup Service (OLS).

### New entities added to PSI-MI CV

#### Entity types

Entities to be represented can encompass different kinds of molecules, such as proteins, DNA, RNA, chemicals, drugs, but also non-molecule entities, such as stimuli, phenotypes and biological processes which are fundamental to understanding and representing signalling events. The PSI-MI CV supports the representation of molecular entities (see MI:0313 and children), but not of non-molecules interactors. To fill this gap, we created a ‘causal interactor type’ term (MI:2259) and related children (‘phenotype’ and stimulus’).

#### Biological Role (Directionality)

Causal interactions are directional and therefore, by definition, asymmetric as the interacting pair of entities have two distinguishable roles: one entity constitutes the modulator, while the other one is the target of the modulation. Some terms in the PSI-MI CV already describe such asymmetry, for instance terms like ‘enzyme’ (MI:0501) or ‘enzyme target’ (MI:0502). To generalize this concept, we defined two new ‘biological role’ parent terms: ‘regulator’ (MI:2274) and ‘regulator target’ (MI:2275).

#### Causality (Effect)

A fundamental piece of information required for causality is the definition of the effect (positive or negative) that the regulator entity has on the regulated. Such information was completely missing in the PSI-MI CV. To address this issue, we created the ‘causal statement’ term. Under the causal statement parent term, it is now possible to find all the necessary definitions to explain the biological effect that (the function of) an entity has on (the function of) another entity and therefore provide more details on how a modulator entity acts on a modulated entity. A modulator entity can act by up-regulating or down-regulating (the function of) another entity. The new terms also enable the users to distinguish between the types of regulation and to specify whether the regulation of the modulator entity acts on the activity or the quantity (by controlling the expression or the stability) of the modulated entity.

#### Interaction Type (mechanism)

The major goal of defining terms for the CausalTAB is to enable representation of interactions occurring in signalling events. Many resources, such as SIGNOR, IntAct, and GO, capture data on molecular events underlying causal interactions. In a biological context, signalling events are mostly cascades of physical interactions, enzymatic reactions and post-translational modifications.

The PSI-MI CV already contained all the interaction type terms necessary to represent ‘direct causal interaction’ such as molecular interactions and enzymatic reactions between the partners that is the regulator entity being immediately upstream to the target one (see also Interaction type branch in the PSI-MI CV).

However, the information reported in the literature about the molecular mechanism through which a molecule or an environmental condition influences the status of a downstream entity can be limited. SIGNOR and Signalink also annotate causal interaction where the regulator is not immediately upstream to the target. For example, the information that DNA damage can induce intermolecular autophosphorylation of the protein ATM, through the up-regulation of its kinase activity (Bakkenist & Kastan, 2003).

To enable the representation of this data and to ensure the distinction between ‘direct causal interactions’ and ‘indirect causal interactions’, we created a new interaction type: ‘Functional association’ defined as ‘*Binary relationship between biological entities when one of them modulates the other in terms of function, expression, degradation or stability of the other and the relationship between the partners cannot be ascertained as direct, so intermediate steps are implicitly present. This relation specifically does not imply a physical interaction between the entities involved*’.

To add further information on ‘Functional associations’, we created the ‘causal regulatory mechanism’ that describes: ‘*Type of relationship between entities involved in a causal interaction. This term is to be used only to describe the effect of a modulator entity A on a modulated entity B when A is not immediately upstream of B*’.

To conclude, the result of this revision-integration effort is the creation of 29 new terms, which allow the description of signalling relationships reported in all the repositories mentioned above, using the PSI-MI schema. The full set of terms is available at https://www.ebi.ac.uk/ols/ontologies/mi/terms?iri=http%3A%2F%2Fpurl.obolibrary.org%2Fobo%2FMI_2233.

### The new PSI MITAB2.8 version or CausalTAB

The CausalTAB derives from the PSI-MI standard, enhanced to capture causal interactions among biological entities. HUPO-PSI MITAB2.7 has been extended by adding four more columns allowing the description of signalling data and, therefore, updated to the new PSI MITAB2.8 version or CausalTAB.

CausalTAB maintains the same structure as previous versions of HUPO-PSI MITAB: it is a tab-delimited file where each row corresponds to an interaction. The file is organized in two main parts: one part reports annotations on the interactors while the second part reports annotations on the occurring interaction. The extended HUPO-PSI MITAB format now has 46 columns instead of 42 (fig. 2). This ensures a compatible merge of non-directional binary interactions with both directional and causality data, when available.

**Figure 2:**
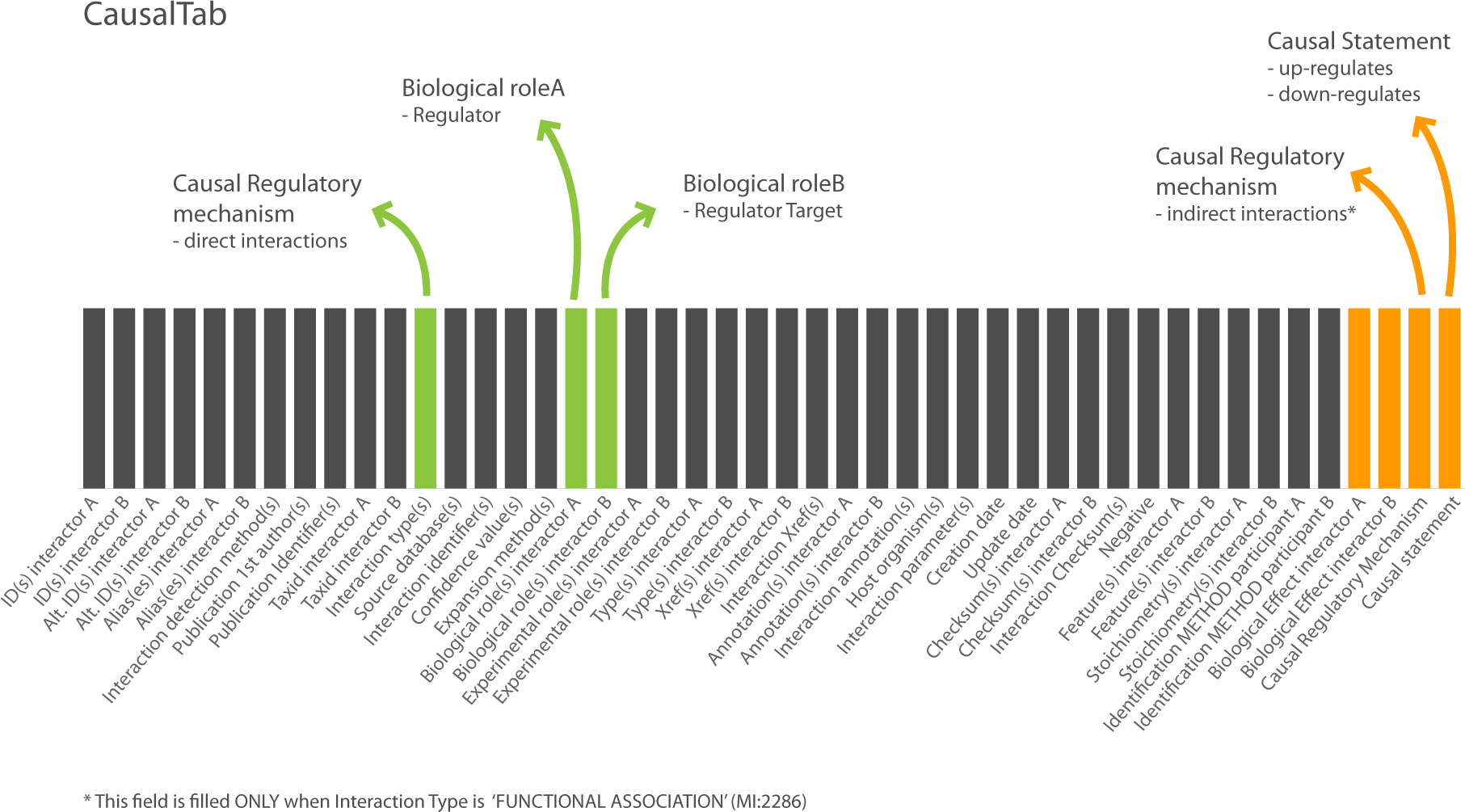
A schematic representation of the CausalTAB. The last four columns (in orange) represent the new columns introduced in the new updated PSI-MI TAB 2.8 version. The arrows indicate those columns of the PSI-MI TAB necessary to describe a causal interaction.

The PSI MITAB 2.7 already contained many columns required to define the elements of a causal interaction, for example: ‘Biological role(s) interactor A or B’, ‘Feature(s) interactor A or B’, ‘Interactor type’.

In particular, the ‘Interaction type’ column will contain all the terms that describe the type of relationship between entities involved in a causal interaction. Terms such as ‘phosphorylation reaction’ (MI:0217) or ‘physical association’ (MI:0915) or ‘ubiquitination reaction’ (MI:0220) are already present in the MI-CV, under the ‘Interaction type’ CV terms and depict all the mechanism underlying a ‘direct causal interaction’. ‘Indirect causal Interactions’ are identifiable by the new interaction type: ‘Functional association’.

The four new columns inserted allow a full export of causal data (fig. 2) and refer to the new controlled vocabulary terms described in the previous paragraph. We, here, briefly discuss the new columns:

The ‘Causal Statement’ column is designed to report the effect of modulator entity A on a modulated entity B. It might contain any child term of ‘causal statement’, including new terms such as ‘up-regulates activity’ (MI:2236) that define ‘the effect of a modulator entity A on a modulated entity B that increases the frequency, rate or extent of the molecular function of B, an elemental biological activity occurring at the molecular level, such as catalysis or binding’.

The ‘Biological Effect Interactor A’ column contains the GO term associated with the Molecular Function of interactor A that is responsible for its regulatory activity. This column will have a GO xref associated. As examples, this column will contain the ‘kinase activity’ GO term, for a kinase phosphorylating its substrate; or the ‘RNA polymerase II transcription factor activity, sequence-specific DNA binding’ GO term for a transcription factor binding the promoter sequence of its target gene.

Similarly, the ‘Biological Effect Interactor B’ column contains the GO term associated with the Molecular Function of interactor B that results to be modulated by the Entity A.

The ‘Causal Regulatory Mechanism’ contains terms that describe indirect causal interactions, such as ‘post transcriptional regulation’, ‘transcriptional regulation’ and ‘translation regulation’, where the effect of entity A is not necessary immediately upstream the entity B. These terms have to be always associated to the term ‘Functional Association’ at the ‘Interaction Type’ level.

The new PSI MITAB2.8, under the name CausalTAB (supplementary fig. 1), is already downloadable through the SIGNOR database under the download page (https://signor.uniroma2.it/downloads.php) and will soon also become a download option through the PSICQUIC web service (Aranda *et al*, 2011), when the new implementation version is released (see Tools section).

### Tools

Over the years, several tools have been developed and maintained in order to empower the PSI-MI format usage. These tools allow, for example, graphical network representation, data format conversions, and use of the schema validation, and are all accessible through the PSI-MI web pages (http://www.psidev.info/groups/molecular-interactions). The new HUPO-PSI MITAB2.8 format is now compatible with most of these tools. The PSI Common QUery InterfaCe (PSICQUIC) web service (Aranda *et al*, 2011) (del-Toro *et al*, 2013), an application that allows the retrieval of PSI-MI standardized interaction data, has recently been updated to be compatible with the new MITAB2.8 format. The Molecular Interactions Query Language (MIQL), a common way to access the data in PSICQUIC and perform search queries has also been updated to the version 2.8 to allow querying the 4 new columns that describe causality. When the new PSICQUIC version will be released, data providers will be able to add causality information and users to download it in the CausalTAB format. The existing PSI MITAB2.7 format documentation is available at the PSICQUIC github page (https://psicquic.github.io/MITAB27Format.html) while the new documentation encompassing the MITAB2.8 format will soon be released and become available through the same Github resource.

Moreover, common methods for interpreting omics data such as gene ontology based functional classification or Gene Set Enrichment Analysis (Subramanian *et al*, 2005) work well for deciphering general areas of biology altered in response to a stimulus, but cannot model the signalling response or uncover the mechanisms of action. This requires causal network analysis methods, and while suitable algorithms are freely available (Bradley & Barrett, 2017), publicly available causal interaction data needs to increase to encourage wider uptake and application for these more recently developed methods.

## Conclusions

There is a strong drive from the scientific community towards the adoption of the FAIR (Findable, Accessible, Interoperable, Reusable) Data Principles that explain which rules data resources, tools and vocabularies should adopt in order to promote data availability and reusability by other users (Wilkinson *et al*, 2016).

Our work has been inspired by the FAIR principles and stimulated by the need to make biological signalling data from disparate resources compatible with each other and consequently interoperable and available to the scientific community.

The signalling community has discussed the new standards during the Causal Reasoning Workshop, the Seattle (2015), Ghent (2016), Beijing (2017) and Heidelberg (2018) HUPO-PSI meetings (http://www.psidev.info/content/hupo-psi-meeting-2017), the Malta and Lisbon COST Action 15205 GREEKC (http://greekc.org/) workshops and in Rome during an ELIXIR-IT funded Curation Workshop on Molecular and Causal Interactions.

During all these meetings, it was possible to revise, discuss and formalise the CausalTab and the new CV terms associated with it and the HUPO-PSI MITAB2.8 update. The proposed format represents the consensus view of the signaling community and it has also been discussed with potential user groups, for example members of the EMBL-EBI Industry program at workshops in Boston and San Diego, US.

Moreover, during the curation workshop, we started to develop new curation rules in order to make the work done for CausalTAB more efficient and useful. Those rules have been accepted by the IMEx Consortium (Orchard *et al*, 2012) and SignaLink curators and will be adopted for the annotation of signalling data.

In the future, we aim to define the fundamental and mandatory information to capture in a causal interaction (i.e, specific information for causal interaction and entity objects, recommendation on ontology to use), by building a guideline called the “Minimum Information about a Causal Statement”, to unify the representation of causal interactions.

Moreover, we will aim at mapping our new terms with the Relations Ontology terms that also contain some terms suitable for the description of causal relations and their directionality.

Thanks to this community effort, we are now able to capture causal interaction data in a structured format, compatible with PSI-MI data and tools. The development of new PSI-MI Controlled Vocabulary terms specific to described and defined signalling events has allowed the update of the HUPO-PSI MITAB to the new 2.8 version that contains all the for the appropriate description of causal interaction. Causal data can be now annotated in a structured format, exchanged and analysed by the users. Moreover, thanks to its structure, the PSI-MI CV can be modified and updated following the community and user needs.

## Acknowledgements

LL was funded by ELIXIR-IIB, the Italian Node of the European ELIXIR infrastructure, LP by Italian Association for Cancer Research [triennial fellowship Starwood Hotels & Resorts id. 18137], MK and JZ by CA15205 Gene Regulation Ensemble Effort for the Knowledge Commons, GC and LP by the DEPTH project of the European Research Council (grant agreement 322749) and by the AIRC (project IG 2017 Id.20322), AL and MA by the Research Council of Norway (project 247727), VT by the Norwegian University of Science and Technology’s Strategic Research Area "NTNU Health", VT and JZ by the ERACoSysMed grant COLOSYS, RCL by the British Heart Foundation (RG/13/5/30112) and the National Institute for Health Research University College London Hospitals Biomedical Research Centre. The IntAct database and EMBL-EBI-based authors received funding from EMBL core funding and Open Targets (grant agreement OTAR-044). TK is supported by a fellowship in computational biology at Earlham Institute (Norwich, UK) in partnership with the Quadrams Institute (Norwich, UK), and strategically supported by Biological Sciences Research Council (grants BB/J004529/1 and BB/P016774/1).

## Supplementary materials

**Supplementary Table 1:** The CausalTAB. The new MITAB2.8 downloaded from SIGNOR database download page. It is a tab-delimited file where each row defines an interaction.

**Supplementary Table 2:** Table that lists all the new terms adopted and introduced in the PSI-MI controlled vocabularies.

## Bibliography

Aranda B, Blankenburg H, Kerrien S, Brinkman FSL, Ceol A, Chautard E, Dana JM, De Las Rivas J, Dumousseau M, Galeota E, Gaulton A, Goll J, Hancock REW, Isserlin R, Jimenez RC, Kerssemakers J, Khadake J, Lynn DJ, Michaut M, O’Kelly G, et al (2011) PSICQUIC and PSISCORE: accessing and scoring molecular interactions. Nat. Methods 8: 528–9

Ashburner M, Ball CA, Blake JA, Botstein D, Butler H, Cherry JM, Davis AP, Dolinski K, Dwight SS, Eppig JT, Harris MA, Hill DP, Issel-Tarver L, Kasarskis A, Lewis S, Matese JC, Richardson JE, Ringwald M, Rubin GM & Sherlock G (2000) Gene Ontology: tool for the unification of biology. Nat. Genet. 25: 25–29

Bakkenist CJ & Kastan MB (2003) DNA damage activates ATM through intermolecular autophosphorylation and dimer dissociation. Nature 421: 499–506

Barabási A-L & Oltvai ZN (2004) Network biology: understanding the cell’s functional organization. Nat. Rev. Genet. 5: 101–113 Available at: http://www.nature.com/doifinder/10.1038/nrg1272

Bradley G & Barrett SJ (2017) CausalR: Extracting mechanistic sense from genome scale data. Bioinformatics 33: 3670–3672

del-Toro N, Dumousseau M, Orchard S, Jimenez RC, Galeota E, Launay G, Goll J, Breuer K, Ono K, Salwinski L & Hermjakob H (2013) A new reference implementation of the PSICQUIC web service. Nucleic Acids Res. 41: W601–6

Demir E, Cary MP, Paley S, Fukuda K, Lemer C, Vastrik I, Wu G, D’Eustachio P, Schaefer C, Luciano J, Schacherer F, Martinez-Flores I, Hu Z, Jimenez-Jacinto V, Joshi-Tope G, Kandasamy K, Lopez-Fuentes AC, Mi H, Pichler E, Rodchenkov I, et al (2010) The BioPAX community standard for pathway data sharing. Nat. Biotechnol. 28: 935–942

Deutsch EW, Orchard S, Binz PA, Bittremieux W, Eisenacher M, Hermjakob H, Kawano S, Lam H, Mayer G, Menschaert G, Perez-Riverol Y, Salek RM, Tabb DL, Tenzer S, Vizcaíno JA, Walzer M & Jones AR (2017) Proteomics Standards Initiative: Fifteen Years of Progress and Future Work. J. Proteome Res.

Fazekas D, Koltai M, Türei D, Módos D, Pálfy M, Dúl Z, Zsákai L, Szalay-Bekő M, Lenti K, Farkas IJ, Vellai T, Csermely P & Korcsmáros T (2013) SignaLink 2 – a signaling pathway resource with multi-layered regulatory networks. BMC Syst. Biol. 7: 7

Hermjakob H, Montecchi-Palazzi L, Bader G, Wojcik J, Salwinski L, Ceol A, Moore S, Orchard S, Sarkans U, Von Mering C, Roechert B, Poux S, Jung E, Mersch H, Kersey P, Lappe M, Li Y, Zeng R, Rana D, Nikolski M, et al (2004) The HUPO PSI’s Molecular Interaction format - A community standard for the representation of protein interaction data. Nat. Biotechnol.

Hucka M, Finney A, Sauro HM, Bolouri H, Doyle JC, Kitano H, Arkin AP, Bornstein BJ, Bray D, Cornish-Bowden A, Cuellar AA, Dronov S, Gilles ED, Ginkel M, Gor V, Goryanin II, Hedley WJ, Hodgman TC, Hofmeyr JH, Hunter PJ, et al (2003) The systems biology markup language (SBML): A medium for representation and exchange of biochemical network models. Bioinformatics 19: 524–531

Kanehisa M, Furumichi M, Tanabe M, Sato Y & Morishima K (2017) KEGG: New perspectives on genomes, pathways, diseases and drugs. Nucleic Acids Res. 45: D353–D361

Kerrien S, Orchard S, Montecchi-Palazzi L, Aranda B, Quinn AF, Vinod N, Bader GD, Xenarios I, Wojcik J, Sherman D, Tyers M, Salama JJ, Moore S, Ceol A, Chatr-Aryamontri A, Oesterheld M, Stümpflen V, Salwinski L, Nerothin J, Cerami E, et al (2007) Broadening the horizon - Level 2.5 of the HUPO-PSI format for molecular interactions. BMC Biol.

Lee MJ & Yaffe MB (2016) Protein regulation in signal transduction. Cold Spring Harb. Perspect. Biol. 8:

Orchard S, Ammari M, Aranda B, Breuza L, Briganti L, Broackes-Carter F, Campbell NH, Chavali G, Chen C, Del-Toro N, Duesbury M, Dumousseau M, Galeota E, Hinz U, Iannuccelli M, Jagannathan S, Jimenez R, Khadake J, Lagreid A, Licata L, et al (2014) The MIntAct project - IntAct as a common curation platform for 11 molecular interaction databases. Nucleic Acids Res. 42:

Orchard S, Kerrien S, Abbani S, Aranda B, Bhate J, Bidwell S, Bridge A, Briganti L, Brinkman FSL, Brinkman F, Cesareni G, Chatr-aryamontri A, Chautard E, Chen C, Dumousseau M, Goll J, Hancock REW, Hancock R, Hannick LI, Jurisica I, et al (2012) Protein interaction data curation: the International Molecular Exchange (IMEx) consortium. Nat. Methods 9: 345–50

Orchard S, Salwinski L, Kerrien S, Montecchi-Palazzi L, Oesterheld M, Stümpflen V, Ceol A, Chatr-aryamontri A, Armstrong J, Woollard P, Salama JJ, Moore S, Wojcik J, Bader GD, Vidal M, Cusick ME, Gerstein M, Gavin A-C, Superti-Furga G, Greenblatt J, et al (2007) The minimum information required for reporting a molecular interaction experiment (MIMIx). Nat. Biotechnol. 25: 894–8

Perfetto L, Briganti L, Calderone A, Perpetuini AC, Iannuccelli M, Langone F, Licata L, Marinkovic M, Mattioni A, Pavlidou T, Peluso D, Petrilli LL, Pirrò S, Posca D, Santonico E, Silvestri A, Spada F, Castagnoli L & Cesareni G (2016) SIGNOR: a database of causal relationships between biological entities. Nucleic Acids Res. 44: D548–D554

Sidiropoulos K, Viteri G, Sevilla C, Jupe S, Webber M, Orlic-Milacic M, Jassal B, May B, Shamovsky V, Duenas C, Rothfels K, Matthews L, Song H, Stein L, Haw R, D’Eustachio P, Ping P, Hermjakob H & Fabregat A (2017) Reactome enhanced pathway visualization. Bioinformatics

Sivade Dumousseau M, Alonso-López D, Ammari M, Bradley G, Campbell NH, Ceol A, Cesareni G, Combe C, De Las Rivas J, del-Toro N, Heimbach J, Hermjakob H, Jurisica I, Koch M, Licata L, Lovering RC, Lynn DJ, Meldal BHM, Micklem G, Panni S, et al (2018) Encompassing new use cases - Level 3.0 of the HUPO-PSI format for molecular interactions. BMC Bioinformatics 19:

Smith B, Ceusters W, Klagges B, Köhler J, Kumar A, Lomax J, Mungall C, Neuhaus F, Rector AL & Rosse C (2005) Relations in biomedical ontologies. Genome Biol. 6: R46

Smoot ME, Ono K, Ruscheinski J, Wang PL & Ideker T (2011) Cytoscape 2.8: New features for data integration and network visualization. Bioinformatics 27: 431–432

Subramanian A, Tamayo P, Mootha VK, Mukherjee S, Ebert BL, Gillette MA, Paulovich A, Pomeroy SL, Golub TR, Lander ES, Mesirov JP, Subramaniana A, Tamayoa P, Moothaa VK, Mukherjeed S, Ebertae BL, Gilletteaf MA, Paulovichg A, Pomeroyh SL, Goluba TR, et al (2005) Gene Set Enrichment Analysis: A Knowledge-Based Approach for Interpreting Genome-Wide Expression Profiles Gene set enrichment analysis: A knowledge-based approach for interpreting genome-wide expression profiles. Source Proc. Natl. Acad. Sci. United States Am. 102: 15545–15550

Türei D, Korcsmáros T & Saez-Rodriguez J (2016) OmniPath: Guidelines and gateway for literature-curated signaling pathway resources. Nat. Methods 13: 966–967

Wilkinson MD, Dumontier M, Aalbersberg IjJ, Appleton G, Axton M, Baak A, Blomberg N, Boiten J-W, da Silva Santos LB, Bourne PE, Bouwman J, Brookes AJ, Clark T, Crosas M, Dillo I, Dumon O, Edmunds S, Evelo CT, Finkers R, Gonzalez-Beltran A, et al (2016) The FAIR Guiding Principles for scientific data management and stewardship. Sci. Data 3: 160018

